# When Maturation is Not Linear: Brain Oscillatory Activity in the Process of Aging as Measured by Electrophysiology

**DOI:** 10.1101/2023.07.26.550635

**Authors:** Sandra Doval, David López-Sanz, Ricardo Bruña, Pablo Cuesta, Luis Antón-Toro, Ignacio Taguas, Lucía Torres-Simón, Brenda Chino, Fernando Maestú

**Affiliations:** Center for Cognitive and Computational Neuroscience, Universidad Complutense de Madrid, 28015, Madrid, Spain; Department of Experimental Psychology, Cognitive Processes and Speech Therapy, Universidad Complutense de Madrid, 28223, Madrid, Spain; Department of Radiology, Universidad Complutense de Madrid, 28040, Madrid, Spain; Department of Psychology, University Camilo José Cela (UCJC), Madrid, Spain; Instituto de Investigación Sanitaria San Carlos (IdISSC), 28040, Madrid, Spain

**Keywords:** *Oscillatory activity*, *Cognitive Neuroscience*, *Magnetoencephalography*, *Maturation*

## Abstract

Changes in brain oscillatory activity are commonly used as biomarkers both in cognitive neuroscience and in neuropsychiatric conditions. However, little is known about how its profile changes across maturation. Here we use regression models to characterize magnetoencephalography power changes within classical frequency bands in a sample of 792 healthy participants, covering the range 13 to 80 years old.

Our results reveal complex, non-linear trajectories of power changes that challenge the linear model traditionally reported. Moreover, these trajectories also exhibit variations across cortical regions. Remarkably, we observed that increases in slow wave activity are associated with a better cognitive performance across the lifespan, as well as with larger gray matter volume for elderlies, while fast wave activity decreases with adulthood.

These results suggest that elevated power in low-frequency resting-state activity during aging may reflect a proxy for deterioration, rather than serving as a compensatory mechanism, as usually interpreted. In addition, it enhances our comprehension of both neurodevelopment and the aging process by highlighting the complexity and regional specificity of changes in brain rhythms. Furthermore, our findings have potential implications for understanding cognitive performance and structural integrity.

## Introduction

Ongoing brain oscillatory activity underlies multiple processes of key relevance for human behaviour, such as long-distance brain networks coordination, cognitive and perceptual processing, plasticity mechanisms, and brain maintenance along life, as has been consistently shown by the literature (Pomper et al., 2021; Rempe et al., 2023; Roux et al., 2012; Sauseng & Liesefeld H.R., 2020; W Klimesch, 2012).

In this context, one classical approach to study electrophysiological activity is through analyzing its spectral properties, which can reveal biomarkers for different pathologies and cognitive processes (Algin et al., 2020; Pal et al., 2020; Zhang et al., 2021). In this regard, it has been shown that alpha band oscillations play a crucial role in attention and memory processes, suppressing distracting stimuli and influencing working memory performance. These findings suggest that alpha-band oscillations could be targeted for interventions aimed at enhancing cognitive performance (Jensen et al., 2002; Palva et al., 2010). However, very little is known about the typical diachronic evolution of these markers in healthy development, which should represent an important pre-requisite to interpret, for example, the role of alpha power in different experiments covering different age ranges.

Nonetheless, maturation is a complex phenomenon encompassing various biological processes, including hormonal, fungal microbiota, metabolism, and skeletal development, among many other changes (Fiers et al., 2020; Fleming et al., 2012; Forman et al., 2012). Cerebral changes play a crucial role in this process, involving functional, cognitive, and cellular events such as myelination, synaptic pruning, or cell shrinkage at different stages (Galloway, 1993). Furthermore, the maturational trajectory of different cortical regions varies largely, as different areas evolve in different time windows (Rempe et al., 2023; Toga et al., 2006). Particularly, limbic, auditory, and visual cortices myelinate early, while frontal and parietal neocortices continue myelination into adulthood (Coupé et al., 2017; Toga et al., 2006). Longitudinal studies have described how grey and white matter volume vary over the cortex, following different trajectories and tendencies across the lifespan (Huttenlocher & Dabholkar, 1997; Morrison & Hof, 1997; Sowell et al., 2003). In fact, these dynamic changes has been shown to begin in the early stages of brain development (Gibb & Kovalchuk, 2018; Leidal et al., 2018). As brain oscillations rely on interneuronal and interassemblies communication, it is expected that brain rhythms also change throughout life (Schnitzler & Gross, 2005).

Classically, power changes during maturation have been described using simplistic models based on linear trajectories, potentially due to restricted sample sizes and/or methodological constraints. This classical conceptualization assumes, therefore, monotonic and constant changes through the different stages of life, which may not adjust to the developmental complexity observed in other systems, such as sexual maturation, more prominent in puberty, or even body morphology (Manna, 2014; Sherar et al., 2010). In contrast, Rempe and colleagues (Rempe et al., 2023) described power changes across lifespan including both linear and quadratic models, finding that higher order polynomial models might adjust better to maturational brain processes.

It is even more striking that, besides its scarcity and relative methodological simplicity, previous literature is quite inconsistent regarding the direction of their findings, reporting both increases and decreases with age. Specifically, for slow bands, i.e., delta (2-4 Hz) and theta (4-8 Hz), some studies reported an increment in power across life (Breslau et al., 1989; Klimesch, 1999a), whereas others found a linear decrease (Babiloni et al., 2006; Leirer et al., 2011; Vlahou et al., 2014a) or even no significant modulation (Stacey et al., 2021). Similarly, results for the alpha band, which usually dominates ongoing oscillatory activity during wakeful resting state with eyes closed, show in some cases a linear decrease of power with aging, particularly in right parietal areas (Barry & de Blasio, 2017; Scally et al., 2018), while in others the authors were unable to find any significant age-related modulation (Stacey et al., 2021). The trajectory of beta and gamma bands also presents conflicting findings, with some studies reporting increases in beta power (Stacey et al., 2021; Vlahou et al., 2014a), others decreases in gamma (D. V. P. S. Murty et al., 2020), and others reporting no change with age (Pollock et al., 1990; Vlahou et al., 2014b).

On the other hand, the evolution of the individual alpha peak frequency (IAPF), a reliable and heritable trait of the human power spectrum (Posthuma et al., 2001; Steriade et al., 1990), seems to be more consistently described in the literature. Previous works systematically describe an initial increase in the frequency of the peak during adolescence (Bazanova, 2008; Dustman et al., 1993) followed by a stabilization during adulthood and, eventually, a decrease in later years (Chiang et al., 2011).

Both cognitive performance (Cesnaite et al., 2021; V. P. Murty et al., 2009; Salthouse, 2010) and brain structural integrity (Good et al., 2001; Jernigan et al., 2001) represent solid hallmarks of aging and the maturational process (Van Blooijs et al., 2023). Since electrophysiological activity (Jensen et al., 2019) and neuronal/structural integrity (Trammell et al., 2017; Voineskos et al., 2012) are strongly associated with multiple sensory and cognitive processes, it is a reasonable assumption that changes in these processes might be related, at least to some extent, to changes in electrophysiological activity along life. Therefore, studying the relationship between these variables (electrophysiological, structural, and cognitive evolutions) might help unveiling the mechanisms underlying the maturational process.

The primary objective of this study is to characterize, comprehensively, the electrophysiological changes in healthy development that have shown inconsistent findings in previous literature. We anticipate observing a complex, non-linear patterns of changes in oscillatory activity, influenced both by brain region and frequency band. Additionally, we aim to investigate whether these changes in brain activity during maturation are associated with established biomarkers for brain health, such as cognitive performance in specific domains or integrity of grey matter. Understanding these relationships will enable us to determine whether these changes can be interpreted as a scaffolding compensatory mechanism or rather as a sign of brain deterioration during normal aging.

## Materials and Methods

### Experimental design

The aim of this study was to address oscillatory activity changes across the lifespan in an initial sample including 1036 healthy participants, trying different adjustments to fit the evolution of power in the different frequency bands. This study employed a cross-sectional design, including power as the independent variable and age as the dependent variable.

### Subjects

After MEG scan quality assessment, the final sample for the analyses included 792 right-handed and native Spanish speakers (460 females and 332 males), aged between 13 and 80 years old (Mean: 46.54, SD ± 22.32). Figure 1 shows the age and sex distribution of the sample.

**Fig. 1.**
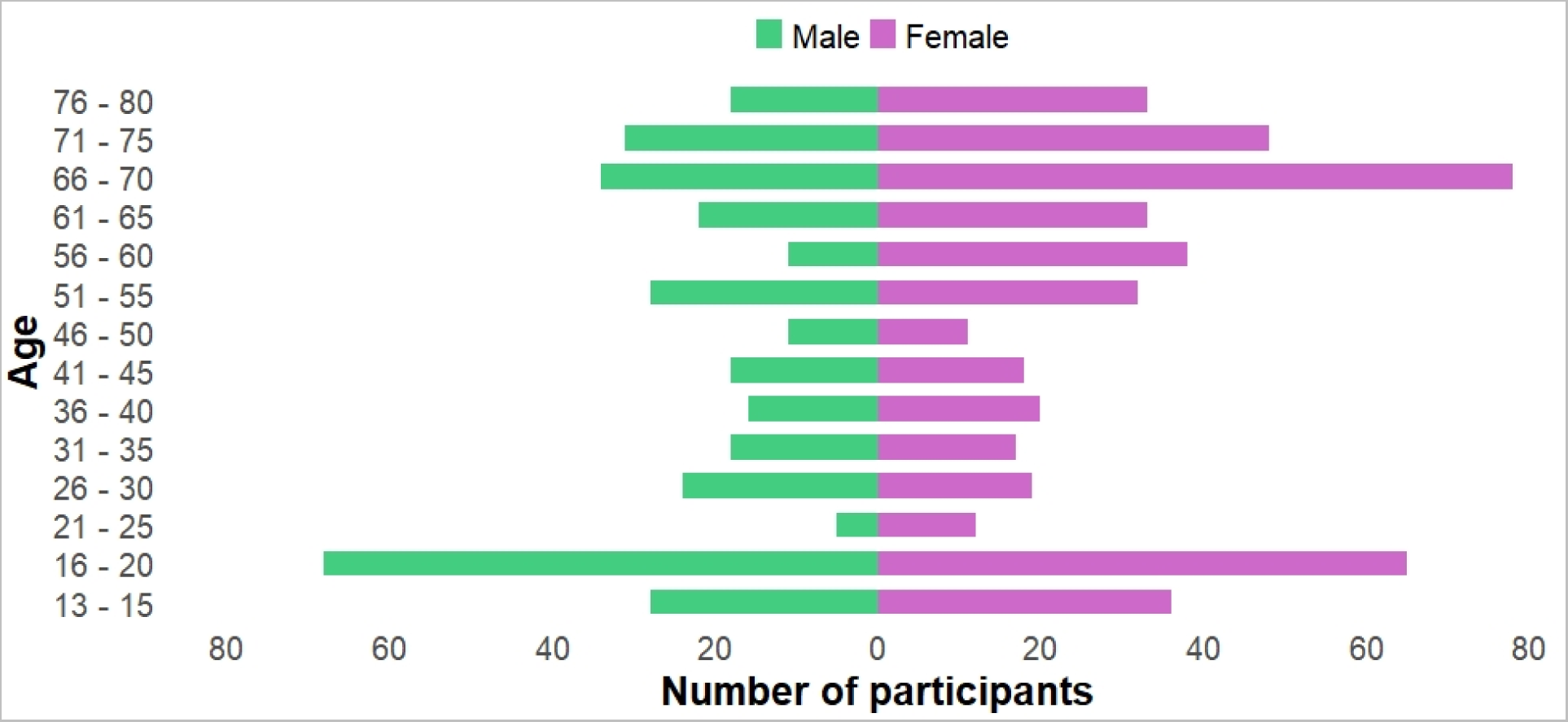
Age-sex distribution of the sample. Figure shows the age and sex distribution of the sample. The x-axis shows the number of participants of each gender (females in purple, males in green) for each age interval represented in the y-axis.

The participants were recruited through different research projects taking place during recent years in the Laboratory of Cognitive and Computational Neuroscience at the Center for Biomedical Technology of Madrid (CTB). Each of these projects was approved by the Ethics Committee of the “Hospital Clínico San Carlos” (Madrid), and all the procedures involved in each project were performed in accordance with the approved guidelines. All the participants included were assessed employing the neuropsychological tools appropriate for each age range and specific project proposals. However, all of them met criteria for being catalogued as healthy volunteers without any cognitive or physical condition that could have interfered with MEG scanning.

### MEG recordings

Electrophysiological data were acquired using a 306 channel Vectorview magnetoencephalography (MEG) system (Elekta AB, Stockholm, Sweden), composed by 102 magnetometers and 204 planar gradiometers. The MEG scanner was placed into a magnetically shielded room (VacuumSchmelze GmbH, Hanau, Germany) at the “Laboratory of Cognitive and Computational Neuroscience” (Madrid, Spain). Recordings were obtained under resting-state condition with eyes-closed, and MEG acquisition consisted of at least 4 minutes of signal for each participant.

Firstly, head shape was obtained by using a three-dimensional Fastrak digitizer (Polhemus, Colchester, Vermont). For each participant, three fiducial points (nasion and left/right preauricular points) and approximately 300 points of the scalp surface were recorded. Furthermore, four head position indication (HPI) coils were placed on their head, two in the forehead and two in the mastoids, and their position was acquired with the same device, allowing head movements to be tracked continuously during MEG recording. To capture eye movements and blinks, a vertical electrooculogram (EOG) was placed on the left eye. In addition, the electrocardiogram (ECG) was obtained from most of the participants by placing a pair of bipolar electrodes at the left mid clavicle and lower right rib bone, respectively.

MEG, EOG, and ECG data were acquired at a 1000 Hz sampling rate using an online anti-alias bandpass filter between 0.1 and 330 Hz.

### MEG pre-processing

Recordings were processed offline using the spatiotemporal signal space separation (tSSS) algorithm (Taulu & Simola, 2006) (correlation window 0.9, time window 10 s) to remove magnetic noise originated outside the head, as well as to correct participants’ head movements during recordings. After spatiotemporal filtering, only magnetometers data were used in the analysis, due to the redundancy with the information contained in the gradiometers (Garcés et al., 2017).

Then, muscular, ocular, and jump artifacts were identified using an automatic procedure from Fieldtrip package (Oostenveld et al., 2011), and visually confirmed by a MEG expert. Afterwards, a blind source separation method (Belouchrani et al., 1997) was employed to remove electrooculogram and electrocardiogram components, when clearly identified. Last, data was segmented into epochs of 4 seconds of continuous, artifact-free data.

### Source reconstruction

Our source model consisted of 2459 sources located in a homogeneous grid of 1 cm defined in the Montreal Neurological Institute (MNI) template. From these 2459 source positions, only those 1202 falling in an area labelled as cortical according to the automated anatomical labeling atlas (AAL) (Tzourio-Mazoyer et al., 2002) were considered. This source model was linearly transformed to the individual head shape obtained prior to the MEG acquisition. The head model consisted of a single shell interface, based on the inner skull surface of the MNI template, and was also linearly transformed to the individual head shape. Last, the lead field matrix was calculated using a modified spherical solution (Nolte, 2003).

MEG data were band-pass filtered between 2-45 Hz using a 450^th^-order finite input response (FIR) filter designed using Hann window. A two-pass was used to filter the data in order to avoid phase distortion. In addition, to avoid edge effects, 2000 samples of real data were added, at each side of the signal’s segment, as padding.

Finally, the activity of each participant in each of the 1202 cortical sources was reconstructed using a linearly-constrained-minimum-variance (LCMV) beamformer (Veen et al., 1997) .

### Spectral Analysis

MEG power spectra were computed using Fieldtrip toolbox and a multi-taper method based on discrete prolate spheroidal sequences, with 1 Hz smoothing, as taper. The spectral power distribution of each epoch was estimated in the range from 2 to 45 Hz in steps of 0.25 Hz, obtaining a vector containing the power for 173 frequency steps for each cortical source. This absolute power was then averaged across epochs and normalized by dividing each value by the total power in the 2-45 Hz range, getting relative power values for each frequency step, source, and participant (173x1202x792).

As the main interest of this work was to describe how the amplitude of classical bands evolves throughout life, we defined of bands of interest according to traditional ranges employed in the literature: delta 2 – 4 Hz, theta 4 – 8 Hz, alpha 8 -12 Hz, beta 20 -30 Hz, and gamma 30-45 Hz. Relative power of each frequency band, source, and subject was estimated as the sum of the relative power steps within each of these ranges. Furthermore, we identified the individual alpha peak frequency (IAPF) using a recently developed algorithm (Donoghue et al., 2020). This algorithm parametrizes the spectrum in the signal of interest by progressively fitting an aperiodic component and adding subsequent putative periodic oscillatory peaks, and one of these oscillatory peaks is used to describe the alpha activity. For our analysis, we estimated de IAPF for each participant using the average power spectrum of those sources labeled as belonging to the occipital lobe, a well-known generator of the alpha rhythm.

### Statical Analysis

#### Relationship between relative power and age

To delve into the evolution of the different brain rhythmś power across the lifespan, we conducted a series of multiple regression analyses in our sample, separately for each of the 1202 cortical sources. A set of regression analyses was performed trying to fit the power in a specific frequency range as a function of the age of each participant. To account for the most plausible different trajectories that power might follow during life, three possible regression analyses were performed, following these equations:

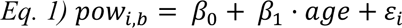

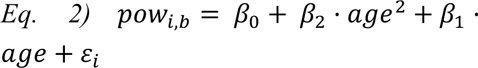

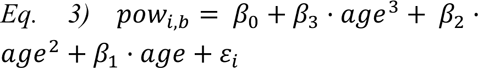

All the models were carried out separately for each cortical source including 792 observations for the two main variables in the models. 𝑝𝑜𝑤_𝑖,𝑏_ contained the relative power for each source (*i*) in each of the bands (*b)* (i.e., delta 2-4 Hz, theta 4-8 Hz, alpha 8-12 Hz, beta 12-30 Hz and gamma 30-45 Hz). Furthermore, an additional set of regressions was carried out for the vector containing the IAPF of each participant. Secondly, the variable 𝑎𝑔𝑒 contained a vector of 792 elements informing the age, in years, of each participant at the time of the MEG signal acquisition.

The first equation intended to explain power change during lifetime as a linear function of age, aiming at capturing monotonic and constant modulations of oscillations’ amplitude from early adolescence (13 years old) to a relatively late age (80 years old). The second model (*Eq. 2*) aimed to explain power modulations across the lifespan including a quadratic term to capture a possible change in the trend of power evolution, from an increase in early stages to a decrease in later years, or vice versa. Lastly, the third regression model (*Eq. 3*) included a cubic term to capture more complex dynamics of power change that could include different trajectories of change during life alternating with periods of relative stability.

Lastly, to decide which model better captured the dynamics of oscillatory activity during life, the goodness of fit of the three possible fittings were statistically compared pairwise by means of a likelihood-ratio test, or Wilks test, selecting the highest order model significantly better compared to the immediately less complex model. All p-values were corrected using false discovery rate (FDR) (Benjamini & Yekutieli, 2005) (Q = 0.05) to account for the multiple statistical tests performed.

#### Relationship between oscillations change and cognitive/brain structure status

Once the main trajectories across life for each frequency band were established, we aimed to find the possible functional implications of these findings. To this aim, quantitative neuropsychology was available for young and order adults.

##### A) Cognitive scores

###### A.1) Young adults

For this age group, the quantitative neuropsychological data base available included the following *BIS-11* subscales, measuring three subtypes of impulsiveness: *cognitive* (attentional) impulsiveness (inattention and cognitive instability)*; motor* impulsiveness (motor disinhibition)*; non-planned* impulsiveness (intolerance of cognitive complexity and lack of self-control) and the *total score* (Patton et al., 1995.). This subset of the sample included a total of 129 participants (67 females and 62 males, 14-20 y/o), with a mean age of 17.18 y/o.

###### A.2) Older adults

This subset of the sample included a total of 331 participants (222 females and 109 males) with available data and meeting age criteria (those over 55 years of age). In order to reduce the dimensionality of the cognitive performance database of older adults, a principal component analysis (PCA) was employed including the scores in the selected tests relevant to five different main cognitive domains that were included in our analyses. 1) *Declarative memory* included the following subtest of the battery Wechsler Memory Scales (WSM-III) (Wechsler, 1997): <u>Logical Memory</u> subtest, which measures verbal memory for stories and is divided into the Immediate and Delayed units; the Memory Logic subtest that assesses the ability to recall information from short paragraphs and is divided into Immediate and Delayed Topics. Additionally, the WMS-III subtest included into this domain were also the Total Recall Word List and the Delayed Recall Word List Topics subtests, which evaluate the ability to recall words from a list immediately and after a delay, respectively.

2) *Working memory* included two subtests from the WSM-III, specifically direct and inverse digit. 3) *Speed processing,* obtained using scores from trail making tests A and B, both time and hits. 4) *Visuospatial memory* included Rey–Osterrieth complex figure neuropsychological assessment, time, and copy accuracy. 5) *Verbal fluency* contained both FAS-phonemic and semantic fluency score.

##### B) Structural integrity scores

Volumetric measurements derived from the MRI scans of each participant were available only for a subset of our sample, corresponding to those projects focused on older adults. Given that most of the controversy in the interpretation of these hypothesized changes in brain activity comes from these later stages, this specific analysis is focused only on the older adult segment of our sample..

T1-weighted MRI images from each participant were produced using a General Electric 1.5T system and applying a high-resolution antenna and a homogenization PURE filter (fast spoiled gradient echo sequence, TR/TE/TI = 11.2/4.2/450 ms; flip angle 12°; 1 mm slice thickness, 256 × 256 matrix, and FOV 25 cm). We used FreeSurfer software (version 6.1.0) for automated cortical parcellation and subcortical segmentation (Fischl et al., 2002). The structures included in the analyses were the total cortical gray matter volume (GM) and the “posterior cingulate plus precunei” (PCC) GM, obtained as the result of adding up the volumes (in mm^3^) of both bilateral posterior cingulate cortices and both precunei, collapsed to get a single measure.

###### General procedure

We conducted a series of partial correlation analyses (ruling out the effect of age) to find out whether a higher or lower power value in a specific frequency range for each participant was associated with a better or worse cognitive and/or brain structural status. This would allow us to infer the possible functional meaning of the changes observed in this stage. More specifically, we conducted a partial correlation in each classical band separately for each cortical source, leading to a total of 1202 correlations in each frequency range and variable included in the analysis. To control the statistical effects of multiple testing, we performed a cluster-based permutation test (Maris & Oostenveld, 2007), grouping those contiguous sources showing a significant correlation. Each partial correlation should meet a certain level of significance to be considered as part of the cluster (α-cluster = 0.01). Lastly, the significance of the cluster was compared to the null-distribution of surrogates as described in the method (Maris & Oostenveld, 2007), to obtain the p-value associated to the magnitude of the cluster (α = 0.05).

## Results

### Oscillatory power changes across the lifespan

Here, we firstly describe the different adjustments used to describe power changes trajectories for all classical frequency bands and the IAPF across lifespan. Table 1 summarizes the coefficients values of each of the presented models for each of the adjustments (linear, quadratic, and cubic) and each frequency band, including the r-squared and p-value of each model.

**Table 1.** Correlation coefficients, r-squared and p-values, are summarized for each frequency band and model (linear, quadratic and cubic). *Int* stands for intercept values, *x*, *x^2^*and *x^3^* represents linear, quadratic and cubic terms respectively. *n.s.* stands for not significant.

|  | Lineal |  |  | Quadratic |  |  | Cubic |  |  |
| --- | --- | --- | --- | --- | --- | --- | --- | --- | --- |
|  | Beta / p-value | r-squared | model p-value | Beta / p-value | r-squared | model p-value | Beta / p-value | r-squared | model p-value |
| **Delta** | n.s. | n.s. | n.s. | n.s. | n.s. | n.s. | int: 0.3642<br>x -0.013/7.6e-26<br>x<sup>2</sup> 0.00024/1.003e-17<br>x<sup>3</sup> -1.57e-06/2.96e-14 | 0.4771 | 1.80E-110 |
| **Theta** | int: 0.240<br>x: -7.693e-04/<br>1.174e-53 | 0.2601 | 1.17E-53 | int: 0.287<br>x: -0.003 / 1.668e-16<br>x<sup>2</sup>: 2.33e-05 / 8.809e-09 | 0.2671 | 5.61E-54 | int: 0.345<br>x -0.01/3.047e-09<br>x<sup>2</sup> 1.97e-04/1.97e-06<br>x<sup>3</sup> -1.21e-06/4.68e-05 | 0.1775 | 3.56E-33 |
| **Alpha** | int:0.167<br>x: 8.98e-04 / 3.306e-32 | 0.162 | 3.31E-32 | int: 0.2189<br>x: -0.0018/7.190e-06<br>x<sup>2</sup>: 1.794e-05/6.471e-05 | 0.0332 | 1.66E-06 | int: 0.2208<br>x p: 0.0106/0.001<br>x<sup>2</sup> -2.773e-04/3.42e-04<br>x<sup>3</sup> 2.0264e-06/2.78e-04 | 0.0324 | 9.61E-06 |
| **Beta** | int: 0.293<br>x: 0.0011 / 3.878e-48 | 0.236 | 3.88E-48 | int: 0.2189<br>x: 0.0054 / 1.373e-23<br>x<sup>2</sup>: -4.02e-05 / 6.734e-12 | 0.373 | 1.08E-80 | int: 0.1307<br>x 0.012/1.2729e-09<br>x<sup>2</sup> -2.054e-04/2.48e-05<br>x<sup>3</sup> 1.123 e-06/0.0013 | 0.3373 | 5.20E-70 |
| **Gamma** | int: 0.0552<br>x: 4.53e-04 / 3.593e-27 | 0.1372 | 3.59E-27 | int: 0.0542<br>x: 0.0011 / 3.953e-06<br>x<sup>2</sup>: -9.4962e-06 / 2.081e-04 | 0.0562 | 1.22E-10 | int: 0.0426<br>x 0.0048/1.23e-08<br>x<sup>2</sup> -9.21e-05/4.061e-06<br>x<sup>3</sup> 5.53e-07/1.16e-04 | 0.1195 | 1.30E-21 |

#### Delta

For delta band, the adjustment that better describes the changes in life continuum across the entire brain surface (as can be seen in Fig.2. panel A) was the cubic (r-squared = 0.4771; p-value = 1.8 e-110). This regression showed, in average, a noticeable decreasing trajectory during the first decades of the continuum (13-30 years old), followed by a stable period with no pronounced changes extending until the age of 55 years old, approximately. From this age on, delta power showed, again, a decreasing trend, to a lesser amount compared to the previous maturational stages. This is, the magnitude of the decrement becomes increasingly pronounced following the age of 55, and although reductions are observed in earlier stages, they are comparatively less steep than those seen within this age cohort.

**Fig. 2.**
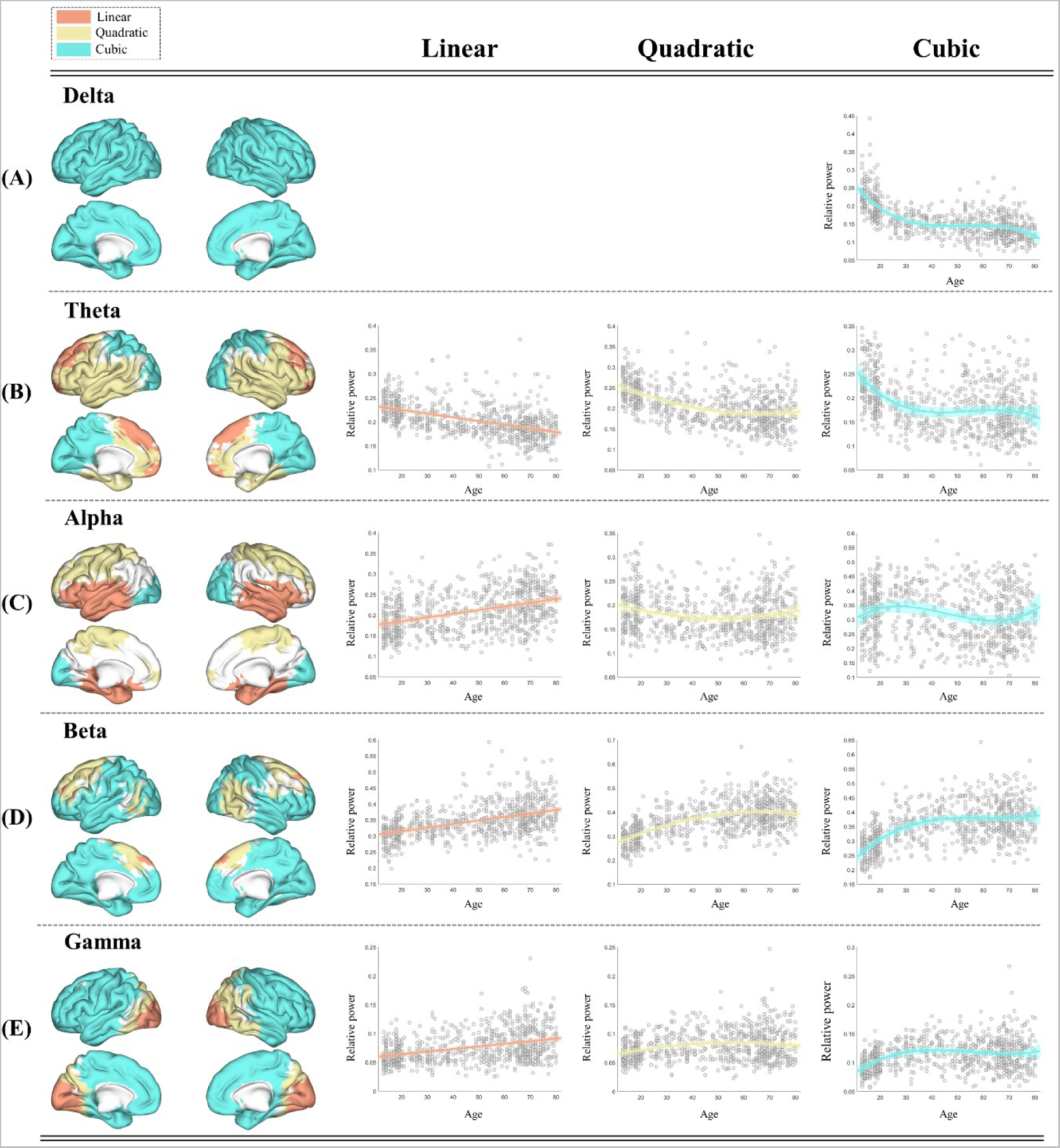
Patterns of power changes throughout life for each classical band. Figure shows for each frequency band (row panels; **A** to **E**) the mean scatter plot of the significant sources for each regression model: linear, quadratic and cubic (columns; left to right). In each scatter plot, relative power is presented in the y-axis and age in years in the x-axis. The different colours on the brain surfaces represent the statistically best adjustment for each source: light orange: linear; yellow: quadratic; light blue: cubic. Confident intervals are represented as a shadow around the tendency line.

#### Theta

For the theta band, a significant model showing linear adjustment with a decreasing tendency (r-squared = 0.2601; p-value = 1.17e-53) was the one better describing the changes in superior frontal areas, specifically over the bilateral frontal gyrus (orbital and medial) and across bilateral supplementary motor areas (Fig. 2, panel B, left section). In addition, a significant U-shaped quadratic model (r-squared = 0.2671; p-value = 5.61e-54) gave a significant better fit to the changes across the lifespan over bilateral temporal structures (hippocampi, temporal poles, and temporal gyri), over ventral frontal areas (inferior opercular and orbital frontal gyrus), in the insula, and in a small portion of the parietal lobe around the Rolandic operculum (Fig. 2, panel B, middle section). These regions showed a pronounced decrement in the first decades of life (13-30 years old), followed by a less intense fall until 55-60 years old, when a change into a positive trend starts. Finally, the cubic adjustment gave a better explanation of the changes (r-squared = 0.1775; p-value = 3.56e-33) over bilateral superior parietal lobes and occipital areas, including the posterior part of the cingulate gyri, the calcarine fissures, cunei, precunei, and lingual gyri (Fig. 2, panel B, right section). Cubic fitting for theta band followed a similar tendency than that for delta band, with a decreasing tendency at the first decades, a plateau during middle age, and an inflexion point that changes the course of the trajectory into a decrease from around 55 years on.

#### Alpha

Changes in alpha band across life maturation were best explained by a linear fitting (r-squared = 0.162; p-value = 3.31e-32) in bilateral temporal areas covering the hippocampi and parahippocampal cortices, inferior frontal gyri, and over temporal poles (Fig. 2, panel C, left section). However, a quadratic adjustment (r-squared = 0.0332; p-value = 1.66e-06), in which alpha power decreases its amplitude from age 13 to 35-40 years old and then increases from 60 years on, was the best model for the sensorimotor cortex, particularly over bilateral paracentral lobe, superior frontal, precentral, and postcentral gyri, and across bilateral supplementary motor areas (Fig. 2, panel C, middle section). Last, in occipital areas a cubic fitting explained power evolution significantly better than simpler models (r-squared = 0.0324; p-value = 9.61e-06). The regions explained by this model included bilateral occipital structures such us calcarine, cunei, and lingual gyri (Fig. 2, panel C, right section). Occipital changes were characterized by a subtle power increment during the first decades included in the analysis, followed by a decrease in middle age, ensued from a noticeable increment at last decades.

#### Beta

The linear fitting (r-squared = 0.236; p-value = 3.88e-48) described the beta band power increase observed throughout life in premotor areas, including a small portion of the medial superior frontal gyrus (Fig. 2, panel D, left section). Furthermore, quadratic adjustment (r-squared = 0.373; p-value = 1.08e-80) gave a significantly better description of them in a reduced portion of the temporal and the parietal lobe (left angular gyrus), and in bilateral supplementary motor areas, following an inverse U-shaped trajectory, characterized by a noticeable increment from 13 to almost middle ages (35-40 years old), and then a plateau with a less pronounced decrement since (Fig. 2, panel D, middle section). Lastly, a cubic adjustment (r-squared = 0.3373; p-value = 5.20e-70) was the best fitting for the changes in bilateral occipital, temporal. and parietal lobes, and in the bilateral frontal lobe including the gyri recti and pre-post-supramarginal gyri (Fig. 2, panel D, right section). First decades were characterized by a sharp increment, followed by a relatively stable period from 30 to 55 years, and a subsequent, less pronounced, increment from this point forward.

#### Gamma

Gamma band showed a spatial pattern in which progressively more complex adjustments better described power changes in maturation following roughly a posterior to anterior brain axis. The linear fit was the best (r-squared = 0.1372; p-value = 3.59e-27) in bilateral occipital zones, specifically in primary visual zones, including calcarine fissures, cunei, and the lingual gyri (Fig. 2, panel E, right section). A quadratic adjustment (r-squared = 0.0563; p-value = 1.22e-10), showing an inverse U-shape in which gamma power increased until age 55 to then start decreasing its power, was significantly better over visual association areas, particularly in right parietal zones, including precuneus and angular gyrus, and a reduced portion of bilateral posterior cingulate gyri (Fig. 2, panel E, middle section). Last, cubic fitting (r-squared = 0.1195; p-value = 1.30e-21) better explained changes in anterior areas, including bilateral frontal, parietal, and temporal lobes, and a small portion of the occipital lobe (Fig. 2, panel E, right section). The changes distribution of this fitting follows a similar tendency to that of beta band, although with a slightly less remarkable increase during the first decades, a stable period in middle age, and a subtle increase during last decades.

#### Alpha Peak

For the alpha peak, a quadratic adjustment, with an inverted U-shape, gave the better explanation of the change trajectory of the frequency throughout life (r-squared = 0.112; p-value = 3.52e-21) (Fig. 3). The coefficients were: Intercept :9.93; x: 0.032 p-value: 0.0003; x^2^: −0.0005 p-value: 3e-07. The maximum average value of the peak was 10.45 Hz and was reached at the age of 32 years old.

**Fig. 3.**
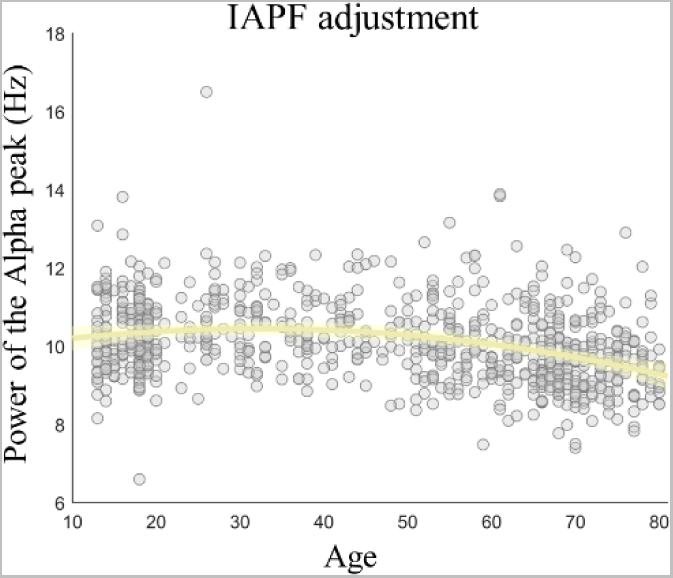
IAPF adjustment throughout life. Figure shows how the IAPF value changes across life span following a quadratic model. The y-axis represents the frequency of the Alpha-peak in Hz and x-axis shows age in years. Confident intervals are represented around the tendency line as a shadow.

### Relationship between oscillations change and cognitive status in younger adults

Here we present the results concerning the relationship between changes in oscillations amplitude in younger adults and their cognitive status. In all cases, the scores of the tests (the Barratt impulsiveness scale, or BIS) represents a higher maturity by lower scores. Therefore, the results of the correlations must be interpreted in reverse, meaning that lower scores indicate a more preserved cognitive status. Again, results will be shown independently for each frequency band.

#### Delta

Delta band showed a significant positive correlation (mean-rho = 0.257; sum-T = 9.3965; p-value = 0.047) between its power and the performance obtained in the BIS cognitive (Fig. 4, panel A, left side, subfigure 1). The significant cluster was mostly located over the right frontal gyrus (including middle, superior, and inferior subregions) and the left gyrus rectus. For non-planned BIS (Fig. 4, panel A, right side, subfigure 2), there was also a positive correlation between performance and power (mean-rho = 0.289; sum-T = 119.3146; p-value = 0.0046), located over bilateral supplementary motor areas and paracentral lobule, cingulate gyrus (anterior and posterior), and left superior frontal gyrus. Lastly, there was a significant positive correlation between the total performance in BIS (Fig. 4, panel A, right side, subfigure 3) and power (mean-rho = 0.271; sum-T = 49.8145; p-value = 0.0122), situated over paracentral lobule, bilateral superior occipital lobe, right superior motor areas, and the right cuneus.

**Fig. 4.**
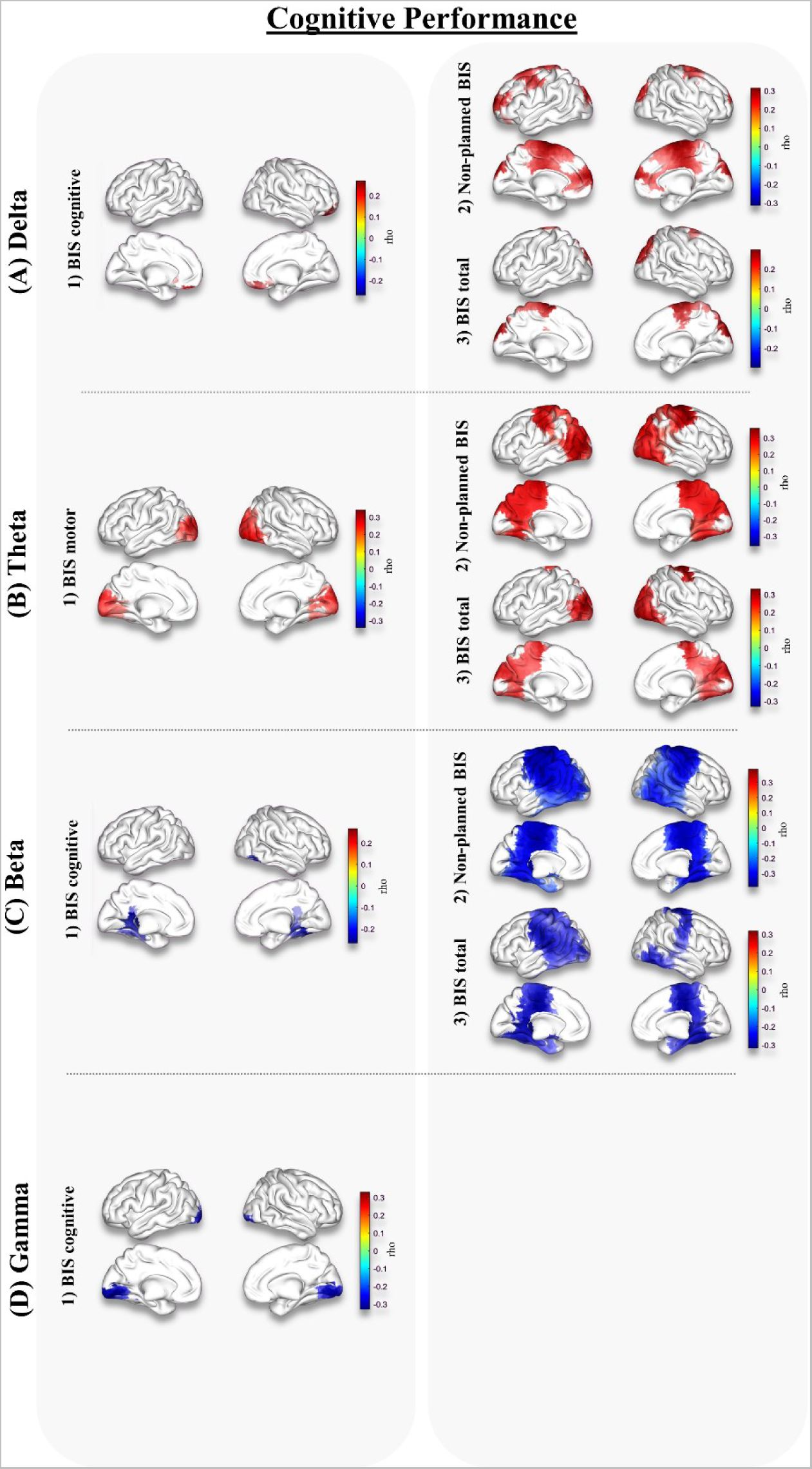
Relationship between power and the cognitive status in the young group. Figure shows the significant correlations between power in each of the classical frequency bands (row panels A to D) and performance in the different subscales of the BIS questionnaire, BIS-cognitive and BIS-motor (left column) or non-planned BIS and total BIS (right column). For each of the brain surfaces colorbar on the right side illustrates rho values of the correlations. Red and blue colours indicate a positive/negative correlation respectively between power and cognition.

#### Theta

Theta band also showed a significant positive correlation (mean-rho = 0.265; sum-T = 51.574; p-value = 0.0143) between its power and the performance in BIS motor (Fig. 4, panel B, left side, subfigure 1). Significant cluster was mostly located over bilateral calcarine fissure, right middle, superior and inferior occipital lobe, and bilateral lingual gyrus. In addition, there was also a significant positive correlation (mean-rho = 0.301; sum-T = 169.3935; p-value = 0.0021) between power in theta and performance in non-planned BIS (Fig. 4, panel B, right side, subfigure 2). This cluster was mostly located over precuneus, superior parietal gyrus, bilateral superior occipital lobe, angular gyrus, cingulate gyrus, and right paracentral lobule. There was also a positive significant correlation (mean-rho = 0.288; sum-T = 105.8750; p-value = 0.0052).

Sources included in the cluster were mostly located over bilateral occipital areas (calcarine and cunei), cingulate gyrus, paracentral lobule, both inferior and superior parts of the left occipital lobe, and the right middle occipital lobe.

#### Alpha

Our findings indicate that there were no significant correlations between alpha band power and scores on any of the subdomains of the Barratt Impulsiveness Scale (BIS).

#### Beta

Beta band power showed a significant negative correlation (mean-rho = −0.253; sum-T= −22.922; p-value = 0.029) with the performance in BIS cognitive (Fig. 4, panel C, left side, subfigure 1). This cluster was located over posterior cingulate gyrus, left lingual gyrus, right fusiform gyrus, and over right inferior occipital lobe. For non-planned BIS there was also a negative correlation (mean-rho =-0.318; sum-T = −227.083; p-value = 0.0006) between power in this band and performance (Fig. 4, panel C, right side, subfigure 2), the significant cluster includes sources located over occipital and parietal lobe, postcentral, precentral, supramarginal and cingulate gyri, Rolandic operculum, hippocampus, precuneus, and supplementary motor areas. In addition, there was also a negative correlation (mean-rho = −0.287; sum-T= −162.901; p-value = 0.0018) between beta and total performance (Fig. 4, panel C, right side, subfigure 3), located over bilateral occipital and parietal areas including paracentral lobe, supramarginal gyrus, cingulate gyrus, postcentral gyrus, fusiform gyrus, and hippocampus.

#### Gamma

For gamma band we found a significant negative correlation (Fig. 4, panel D, subfigure 1) between power and performance in BIS cognitive (mean-rho = −0.299; sum-T = −22.08; p-value = 0.031), located over bilateral calcarine fissures, lingual gyrus, and inferior occipital lobe.

### Relationship between oscillations change and cognitive/brain structure status in aging

Here we show the relationship between changes in oscillations amplitude in older adults and the structural/cognitive status of those participants. Results will be presented separately for each frequency band.

#### Delta

Delta band showed a significant positive correlation (mean-rho = 0.1577; sum-T = 25.2815; p-value = 0.017) between its power and the performance obtained in the visual memory cognitive domain (Fig. 5, panel A, left). The significant cluster was mostly located over the right parietal lobe (angular gyrus, inferior parietal gyrus, and posterior cingulate gyrus) and in the limit between right parietal and frontal lobes (supramarginal gyrus and postcentral gyrus).

**Fig. 5.**
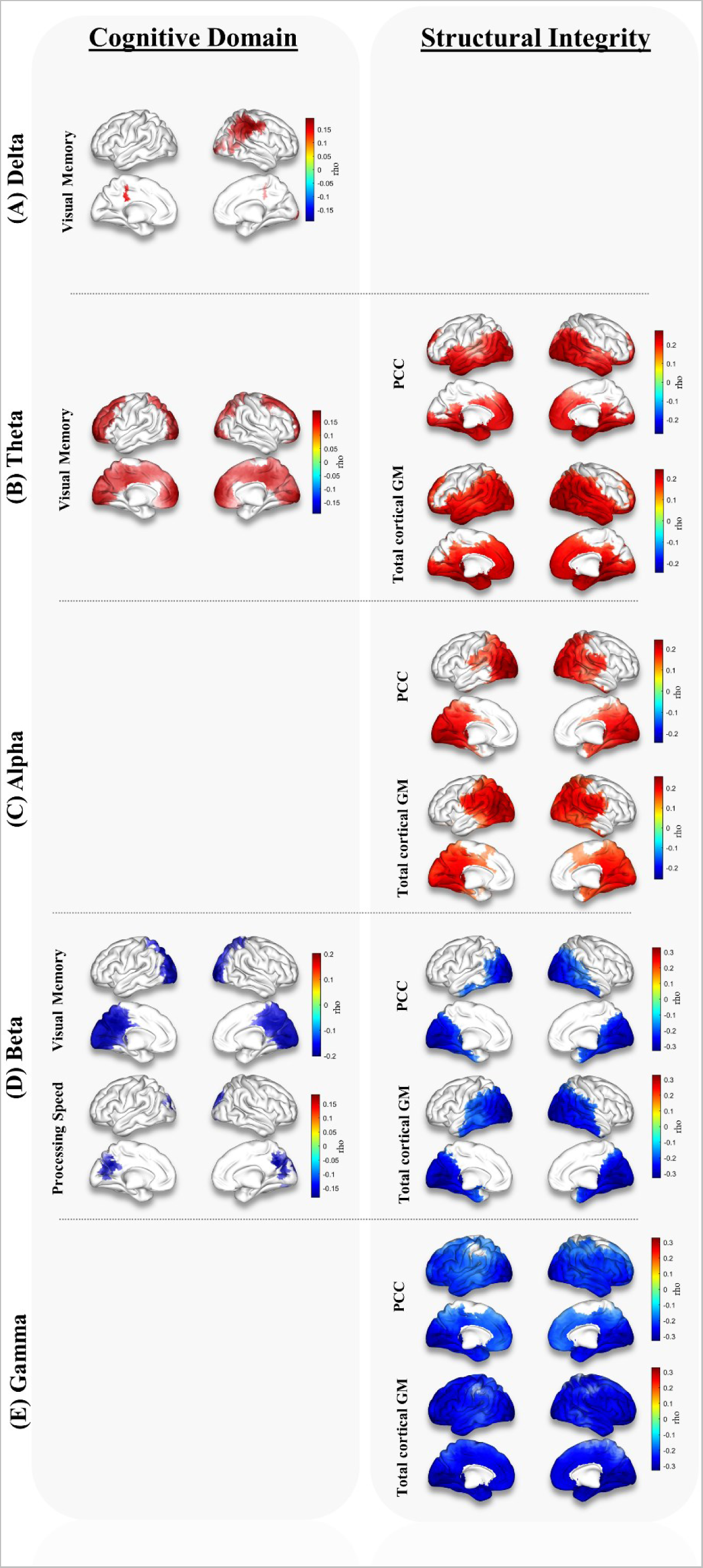
Relationship between power and the structural/cognitive status in older adults. Figure shows the significant correlations between power in each of the classical frequency bands (row panels **A** to **E**) and performance in the different cognitive domains (left column) or structural integrity (right column). For each of the brain surfaces colorbar on the right side illustrates rho values of the correlations. Red and blue colours indicate a positive/negative correlation respectively between power and cognition/structural integrity.

#### Theta

Theta band showed a significant positive correlation (mean-rho = 0.1599; sum-T = 108.8961; p-value = 0.0025) between its power and the performance in visual memory cognitive domain (Fig. 5, panel B, left). Sources included in the cluster were mostly located over bilateral frontal areas (anterior and middle part of cingulate gyri, frontal gyri, and gyri recti), across bilateral parietal structures, and over bilateral occipital areas (calcarine and cunei), covering most of the medial surface of both hemispheres.

Regarding structural integrity (Fig. 5, panel B, right), there was a significant positive correlation between power in this band and the volume of the posterior cingulate cortex (mean-rho = 0.1884; sum-T = 150.9061; p-value = 0.0014), encompassing most of the bilateral temporal lobes (including medial structures such as hippocampi, parahippocampal cortices, fusiform gyri, and amygdalas), and bilateral occipital areas (calcarine, cuneus, and lingual gyrus, among others), and a small portion of the parietal region. In addition, theta band activity over bilateral regions, including most of the temporal and occipital sources, and some frontal (supramarginal gyrus and cingulate gyrus) and parietal regions (inferior parietal gyrus), also showed a significant positive correlation with the total cortical grey matter volume (mean-rho = 0.2101; sum-T= 208.3939; p-value = 1.999e-04).

#### Alpha

No significant correlations between power in the alpha band and cognition were found. Nevertheless, the power of this band was positively correlated with structural integrity (Fig. 5, panel C, right). Specifically, the sources included in the significant cluster (mean-rho = 0.1881; sum-T = 124.5816; p-value = 0.0014) were distributed over posterior cingulate cortex, occipital structures such as the calcarine, cuneus, and lingual gyrus, left Rolandic operculum, bilateral parietal areas including the angular and parietal gyri, and extending over some right hemisphere structures of the temporal lobe (hippocampus, parahippocampal cortices and amygdala). Lastly, alpha power also correlated with total cortical gray matter volume (mean-rho = 0.2009; sum-T = 158.5819; p-value = 3.9996e-04), and the sources of the significant cluster were located over bilateral temporal areas (parahippocampal cortices and hippocampi), bilateral occipital structures (calcarine fissure, cunei, fusiform gyri, and lingual gyri), across bilateral parietal areas (precunei, Rolandic operculum, and posterior cingulate gyri), and over the right temporal gyrus.

#### Beta

Beta band power showed a significant negative correlation (mean-rho = −0.1699; sum-T= −60.369; p-value = 0.0088) with the performance in visual memory (Fig. 5, panel D, left). This cluster was mostly located over bilateral occipital areas (calcarine, lingual gyri, and cunei) and extending into a small portion of the posterior cingulate gyrus and superior parietal lobe. Regarding cognition, there was also a negative significant correlation between beta band oscillatory amplitude and processing speed (mean-rho = −0.1593; sum-T = −23.304; p-value = 0.0226) (Fig. 5, panel D, left), specifically in sources over bilateral occipital areas (cunei and calcarine cortices) and left posterior cingulate cortex.

Furthermore, for structural integrity (Fig. 5, section D, right), beta band power over most of the occipital surface and some bilateral temporal areas including the hippocampus and fusiform gyri, showed a significant negative correlation with the volume of the posterior cingulate cortex (mean-rho = −0.2194; sum-T = −105.3; p-value = 0.0027). Last, there was a negative significant correlation between gray matter volume and beta amplitude (mean-rho = −0.2408; sum-T = −141.0870; p-value = 0.0012), covering most of the bilateral temporal and occipital lobes and, extending into a small portion of the parietal lobule (angular gyrus and posterior cingulate gyrus)

#### Gamma

No significant correlations between the power in gamma band and cognitive performance were found. For structural integrity (Fig. 5, panel E, right), significant negative correlations were found between power in gamma band for both posterior cingulate cortex (mean-rho = −0.21; sum-T = −246.5647; p-value = 9.999e-05) and total cortical grey matter volume (mean-rho = −02576; sum-T = −316.9765; p-value = 9.999e-05). Both clusters occupied most of the cortical surface, being the latter a bit larger and extending more into superior somatosensory areas.

## Discussion

The findings of this study challenge the traditional belief that brain activity follows a linear trajectory over time. Instead, we observed distinct trajectories depending on the frequency and specific cortical brain region. While some brain regions exhibited linear patterns of oscillatory changes across the lifespan, employing more complex adjustments provided a significantly better explanation for the majority of cortical areas. These results support our hypothesis that electrophysiological brain changes have been inadequately characterized in previous research. As a simplified summary, most of the regression results showed that slow bands amplitude decreased as a function of age, whereas for faster rhythms the opposite was true.

Interestingly, our findings contradict the common conception of brain activity slowing with aging (Barry & de Blasio, 2017; Leirer et al., 2011; D. V. P. S. Murty et al., 2020; Scally et al., 2018; Vlahou et al., 2014b). Numerous studies have characterized Alzheimer’s disease (AD) as an accelerated aging process, implying that those changes occurring in AD are those characteristics of the normal aging process, but in an accelerated and pathological way (Franke & Gaser, 2012; Pallas et al., 2008). In fact, multiple studies reported a shift of the power spectrum towards lower frequencies as an electrophysiological hallmark of the disease (Rae Grant et al., 1987) and, thus, this pattern of changes had been commonly associated in the literature to aging in general. Nevertheless, our results showed that, for healthy aging, the opposite pattern was true, meaning that the electrophysiological pattern of changes is qualitatively different in both conditions.

Our results robustly show that the predominant pattern of changes during normal maturation best fitted to a cubic model for several brain regions and frequency bands. We observed that the most significant changes occurred at the extremes of the age range, indicating steeper oscillatory changes in younger and older adults, while the brain activity of middle-aged individuals remained relatively stable.

These patterns align with known brain maturational processes, including myelination/demyelination (Grydeland et al., 2019), plasticity changes, including neurogenesis (Lledo et al., 2006), neuronal pruning (Chechik et al., 1999), and alterations in perineuronal nets and functional networks (Long et al., 2021). Previous studies have reported similar age-related changes with pronounced transformations in early and late stages of life and a relatively stable plateau during middle-ages (Grydeland et al., 2019). Our findings are consistent with these observations, and suggest that the distinct adjustment of oscillatory changes across cortical regions reflects brain developmental events throughout maturation. This comprehensive analysis, conducted with a large and representative sample, contributes to our understanding of brain oscillatory changes, and aligns with findings from other areas of brain maturation research.

The functional implications of aging-related changes have been extensively discussed in the literature (Cabeza et al., 2000; Poulisse et al., 2020); our findings regarding the relationship between oscillatory activity and cognition/structural integrity might shed some light into the current debate about scaffolding and compensation. The scaffolding theory of aging and cognition (STAC) (Park & Reuter-Lorenz, 2009) proposes that increased functional activity in aging serves as a compensatory process, recruiting additional circuits to compensate for age-related declines in cognitive functioning and brain structure. However, our findings do not fully support this view. The correlation analyses indicate that the changes in power spectrum during normal aging represent a sign of brain deterioration, rather than a compensatory attempt, as they are accompanied by worst performance in all the cognitive domains we assessed and a poorer structural integrity.

The evaluation of the relationships between power and cognitive performance helps us to better interpret the neurophysiological development. During early maturational stages, cognitive control plays a critical role in various response processes, including working memory, response selection-inhibition, task-set maintenance, and switching (Badre, 2011; Lenartowicz et al., 2010; Luna et al., 2015); Understanding the dynamics of healthy and impaired cognitive development is of great interest, as cognitive control continues to improve throughout the maturational process, including adulthood. That is why it seems relevant to analyze its relationship with power in early maturational stages, by using scales related to impulsivity as BIS(Patton et al., n.d.), as impulsivity tends to decrease as individuals mature (Caspi et al., 2005; Steinberg, 2008) . For the young group (13-19 y/o), we observed a less pronounced slow wave activity (delta and theta); as higher scores on the impulsivity scale (BIS) are related to an overall worse cognitive status, the positive correlation found between power on these bands and performance in the different subscales of the test would be an indicative of a worsen on cognitive status, reinforcing the characterization of the electrophysiological developmental trajectories we found. Additionally, for this group, a more intense fast wave activity (beta and gamma) negatively correlates with the performance in BIS subscales, reflecting a preserved cognitive and maturational status. Results are coherent with previous findings, in which slow frequency oscillations showed a decrease, and fast waves activity present an enhancement tendency as age increases, in relation with cognitive control processes (Marek et al., 2018).

For the elderly group we observed that a more intense slow wave activity (delta and theta) and a less pronounced fast wave activity (beta and gamma) were associated to overall better cognitive performance. Previous studies have reported a similar positive relationship between cognitive performance and power in slow wave activity (Vlahou et al., 2014b). Finnigan and Robertson (Finnigan & Robertson, 2011) found that higher resting-state theta power at frontal and parietal sites in healthy elderlies was associated with better cognitive functioning, substantiating the important role of slow wave oscillations in neurocognitive function during aging. While slow-wave activity is typically associated with pathology, it has also been shown to dominate during the development of functional networks at early stages of maturation (Feinberg & Campbell, 2010; Marshall et al., 2006). Given its dualistic role, slow wave activity can be interpreted as either a sign of pathological development or a plasticity process.

Numerous studies have shown how increases in power in the theta band in healthy older adults are linked to enhanced cognitive performance, sustained attention, and memory function, specifically in temporal areas, including the hippocampus(Bell et al., 2007; Hsieh & Ranganath, 2013; Klimesch, 1999b). However, similar increases in theta power have also been observed in pathological conditions like the AD continuum, where they are believed to reflect compensatory mechanisms in response to neural damage (Babiloni et al., 2017). Moreover, this enhancement has been also identified as a sign of neuropsychiatric disorder, for example in hyperactivity disorder or depression (Arns et al., 2013). Conversely, Soininen and colleagues (2017) found increased slow-wave activity in both healthy individuals and patients with mild AD following the administration of a food supplement (De Waal et al., 2014).

Therefore, alterations in slow-wave activity do not necessarily are a sign of pathology, but rather a response of the brain to specific interventions or conditions (Karakaş, 2020). Consequently, the interpretation of an increase in slow-wave oscillation must be contextualized within the specific clinical and experimental parameters. In healthy adults, increases at this frequency band might be a marker of healthy neurocognitive aging.

In a similar vein, our results showed that beta oscillatory activity increased during aging, and this was negatively associated with the performance in visual memory and processing speed in last decades. This suggests that a less enhanced beta activity in aging may indicate a better-preserved brain state, which could be reinforced by the findings of Engel and Fries (2010), who concluded that spontaneous enhancement in the oscillatory activity of this frequency band is also associated with impaired movement performance. Regarding the IAPF, no significant correlations were found for cognition or structural integrity. However, our findings are consistent with previous literature that reported an increment of the IAPF during the first developmental stages (20 y/o), with an average mean of the peak value between 9.8-10.5 Hz for young adults, and a progressive slowing from 40 y/o on to 8.5-9.7 Hz(Dustman et al., 1993).

Aging has also been consistently associated with cognitive decline; neurobiological explanations of this decline have focused on grey and white matter loss (Good et al., 2001; Jernigan et al., 2001). Our findings reveal a negative association between grey matter volume and the trajectory of all frequency bands in the last decades. This implies that the decreases observed in theta power and the increases in both beta and gamma rhythms were associated to an overall worse gray matter status. These results, again, reinforce the idea that the electrophysiological changes observed in those decades are a sign of the deterioration accompanying normal aging.

Remarkably, changes in alpha power and cognitive performance were not significantly associated with proxies of brain health in our sample. Alpha activity during resting-state primarily serves the physiological function of inhibiting cortical areas associated with primary processes, as extensively described (Jensen & Mazaheri, 2010; Obleser et al., 2012; van Dijk et al., 2008). Despite sharing a common frequency of oscillation with alpha, classically related to visual cortical areas, two additional rhythms with a dissociate physiological function have been described in this frequency range (Bastarrika-Iriarte & Caballero-Gaudes, 2019; Kuhlman, 1978): the so-called tau (related to audition) and mu (related to somatosensory processing) rhythms. The topographical distribution of these three rhythms in the 8-12 Hz range remarkably aligns with our observed changes. Specifically, a linear trajectory explained changes in the temporal lobe, encompassing the auditory cortex associated with tau-rhythm fluctuations (Lehtelä et al., 1997). Moreover, quadratic adjustments accounted for alpha oscillation changes in sensorimotor cortices where mu-rhythm appears. Lastly, the occipital lobe, housing the main alpha generator and visual cortex, exhibited a cubic trend, with an increment in last decades. This supports the fact that changes in alpha oscillatory activity and its topological distribution across life might be the results of how the different rhythms that fluctuate at this frequency range evolve, following different tendencies in life.

Our study has some limitations that should be acknowledged. Firstly, the cross-sectional nature of the data restricts individual-level conclusions. However, the large number of participants in each age group yields reliable results that help reconcile conflicting literature and expand our understanding through comprehensive lifespan characterization, that as far as we are concerned had not been previously conducted. Secondly, even though at first glance data variability around regression lines is large, a very influential recent work (Marek et al., 2022) demonstrated that as sample size grow, the estimated relationships between neuroimaging and behavioral or demographical variables were much lower than typically reported in the literature, and the replicability of the findings largely increased. Apparently, studies using several hundreds, or even thousands, of participants tend to converge on smaller and more disperse, but more reliable, estimations of any given statistical parameter studied, which seems to fit our relatively variable but highly significant models considering our almost unprecedented sample size in the MEG field. Lastly, data from multiple projects limited the availability of certain information for specific groups (e.g., structural integrity for younger and middle-aged individuals). Future research should replicate these findings longitudinally and investigate the relationship between oscillatory activity patterns and structural integrity in these groups.

As a summary, it is possible to conclude that there is a significant relation between age and the behavior of the different cerebral rhythms. Furthermore, this relation is not strictly linear, and strongly depends on the topographical region as well as the frequency band. Importantly, these changes during late states of aging seem to be related to an overall worse structural integrity and cognition; in contrast, during the early stages, alterations in these oscillations are linked to enhanced cognitive performance.

This characterization analysis will allow to settle a general baseline to compare and assess potential differences between normative and neuropathological electrophysiological data considering its behavior in the different life stages. In addition, it resolves the aging brain scaffolding theory debate, showing that power changes signify deterioration rather than compensation, as they are accompanied by worse cognitive performance and less preserved structural integrity.

These findings can provide, for the first time, a normative data that future studies could use as a reference for cognitive and computational neuroscience, as well as for the characterization of different psychiatric and neurological diseases.

## Conclusions

In conclusion, our study challenges the traditional view of linear brain activity changes across maturation. We found distinct trajectories of brain oscillatory activity depending on frequency and the cortical region. Brain activity challenges prevailing notion of age-related slowing, as oscillatory activity does not simply slow with aging; instead, we observed that slow wave activity decreases while fast wave activity increases. These findings indicate that the electrophysiological pattern of changes in healthy aging is different from pathological conditions. Our results provide valuable insights into neurodevelopment and aging, contributing to our understanding of cognitive performance and structural integrity.

## Funding

Spanish Ministry of Health and Social Politics (National Plan of Drugs); SPI/2010/134 and SPI/2010/051

Ministry of Economy and Competitiveness: General Directorate of Scientific Research; PSI2012-38375-C03-01 and PSI2015-68793-C3-1-R

S.D. and L.T.S. were supported by the predoctoral researchers grant from Universidad Complutense de Madrid and co-founded by Santander bank (CT63/19-CT64/19 y CT42/18-CT43/18 respectively) Ministry of Economy and Competitiveness (PSI2015-68793-C3-1-R, PSI2015-68793-C3-2-R, PSI2015-68793-C3-3-R, RTI2018-098762-B-C31)

Spanish Ministry of Health (PR23/14)

Spanish Ministry of Science and Economy Grants PSI2009-14415-C03-01

## Ethical Approval

All the data collected have been part of projects approved by their corresponding ethics committee within the current legal framework of the Declaration of Helsinki that regulates human experimentation and the guide for good practice in clinical research drawn up by OMS and UNESCO (1995). All the participants signed an informed consent in which they freely expressed their willingness for the data to be analysed within the scope of the research. The processing of all personal data derived from the collection of data to be used in the project complies with the General Data Protection Regulation (RGPD) of the European Union in the acquisition center. In addition, all the neuroimaging and neuropsychology files to be shared are duly anonymized and therefore do not allow any type of traceability between the data and the participant.

## Author contributions

Conceptualization: SD, DLS, FM

Methodology: SD, DLS, PC, RB

Investigation: FM, SD

Visualization: SD, IT, LTS, BD

Supervision: DLS, FM, RB

Fundings: FM

Writing—original draft: SD, DLS, RB, FM

Writing—review & editing: SD, DLS, IT, LTS, BC, LA, PC, RB, FM

## Data and materials availability

The data and codes employed in this manuscript would be shared under reasonable request upon finalization of the project for which they have been gathered.

